# IR spectroscopy: from experimental spectra to high-resolution structural analysis by integrating simulations and machine learning

**DOI:** 10.1101/2025.07.23.665767

**Authors:** Marvin Scherlo, Dominic Phillips, Ricarda Künne, Emiliano Ippoliti, Klaus Gerwert, Carsten Kötting, Paolo Carloni, Antonia S.J.S. Mey, Till Rudack

## Abstract

Understanding biomolecular function at the atomic scale requires detailed insight into the structural changes underlying dynamic processes. Vibrational infrared (IR) spectroscopy—when paired with biomolecular simulations and quantum-chemical calculations—determines bond length variations on the order of 0.01 Å, providing insights into these structutral changes. Here, we address the forward problem in IR spectroscopy: predicting high-accuracy vibrational spectra from known molecular structures identified by biomolecular simulations. Solving this problem lays the groundwork for the inverse problem: inferring structural ensembles directly from experimental IR spectra.

We evaluate two computational approaches, normal mode analysis and Fourier-transformed dipole autocorrelation, against experimental IR spectra of N-Methylacetamide, a prototypical model for peptide bond vibrations. Spectra are derived from simulation models at multiple levels of theory, including hybrid quantum mechanics/molecular mechanics, machine-learned and classical molecular mechanics approaches.

Our results highlight the capabilities and limitations of current theoretical biophysical approaches to decode structural information from experimental vibrational spectroscopy data. These insights underscore the potential of future artificial intelligence (AI)-enhanced models to enable direct IR-based structure determination. For example, resolving the so far experimentally inaccessible structures of toxic oligomers involved in neurodegenerative diseases, enabling improved disease diagnostics and targeted therapies.

**TOC Graphic:** 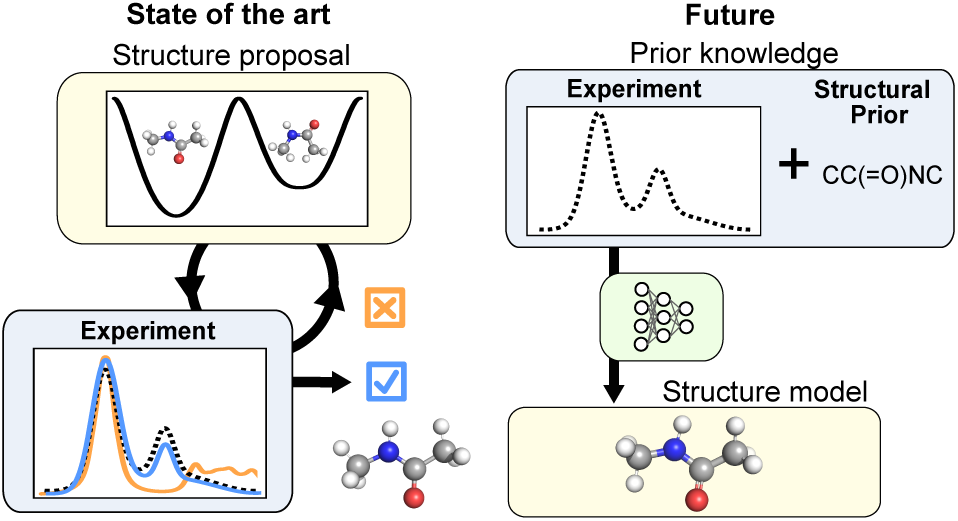

## Introduction

Conformational changes in proteins and biomolecules play a central role in many cellular processes. For many years, spectroscopic methods have been invaluable tools for elucidating the underlying structures and dynamics. While techniques such as Nuclear Magnetic Resonance (NMR) spectroscopy yield atomic-level structures,^1^ and others such as Förster Resonance Energy Transfer (FRET)^2^ and Circular Dichroism (CD)^3^ probe distances and secondary structures respectively, infrared (IR) spectroscopy, stands out for its combination of sensitivity and temporal resolution.^4,5^

The power of IR stems from its ability to resolve minute structural changes, where a ∼ 1 cm*^−^*^1^ frequency shift corresponds to a change in bond length of ∼ 0.001 Å.^5^ While initially used for the qualitative and quantitative analysis of organic compounds, ^6^ IR’s high resolution also makes it ideal for studying enzyme mechanisms and active site dynamics.^7^ By probing the amide I band, IR spectroscopy distinguishes regions of *α*-helices and *β*-sheets, and this has led to its use in biosensors for diagnosing proteinopathies such as Alzheimer’s or Parkinson’s disease.^8,9^ Furthermore, time-resolved Fourier Transform Infrared Spectroscopy (FTIR) offers the unique advantage of being able to capture dynamics on the nanosecond-to-second timescale.^10–12^

However, this remarkable sensitivity creates a significant challenge: the rich information about structure-function relationships is hidden within broad, overlapping vibrational bands that are difficult to assign. Unlike NMR, where spin-spin couplings yield well-defined patterns, IR spectra are more challenging to decode. 2D-IR distinguishes some mode couplings, but it requires femtosecond lasers and suffers from low signal-to-noise ratio. ^13^ Therefore, computational biophysics, especially biomolecular simulations and quantum-mechanical calculations, is essential to translate spectroscopic observables into detailed structural models and mechanistic insights. Historically pioneering work has been done in the context of NMR spectroscopy, bridging experiment and theory, by Prof. Peter A. Kollman and collaborators. In this spirit we dedicate this article. ^14–20^

Historically, computational approaches have focused on solving the *forward problem*: predicting an IR spectrum from a proposed structure (Figure 1A). A human-in-the-loop procedure then iteratively refines the structure until the calculated spectrum matches the experimental one. To obtain a spectrum, two parts are required: a simulation model, which computes trajectories of structural ensembles, and a spectrum model, which computes spectra from the structural data. Depending on the required accuracy, various molecular dynamics (MD) approaches can be employed, such as *ab initio* quantum mechanics (QM), hybrid quantum mechanics / molecular mechanics (QM/MM) simulations, or classical molecular mechanics (MM). Then, to obtain the spectrum, two primary techniques exist. The first, Normal Mode Analysis (NMA), calculates vibrational frequencies from the Hessian matrix of a single equilibrium structure. Warshel’s,^21^ as well as Tavan and Schulten’s work on the retinal structures,^22,23^ validated this approach by obtaining similar IR spectra to experiments. While computationally efficient in its basic form, this approach fails to capture the conformational diversity of molecules at room temperature. This limitation is partly addressed by applying NMA to an ensemble of structures derived from MD simulations either MM or QM/MM.^24–26^ However, this ensemble-based NMA becomes significantly more computationally demanding, as the Hessian matrix must be calculated and diagonalized for a large number of structures. Another approach, typically less computationally demanding, is to compute IR spectra directly from MD simulations via the Fourier transform (FT) of the dipole autocorrelation function,^27–31^ which naturally includes the effects of conformational heterogeneity. However, assigning these IR bands to specific nuclear motions is highly challenging compared to the straightforward approach in normal-mode analysis. Determining isotopic effects requires repeating the entire time-consuming MD simulation, meaning this method typically yields a single, unassigned spectrum, although techniques have been suggested that partially overcome this limitation.^32–34^

**Figure 1:**
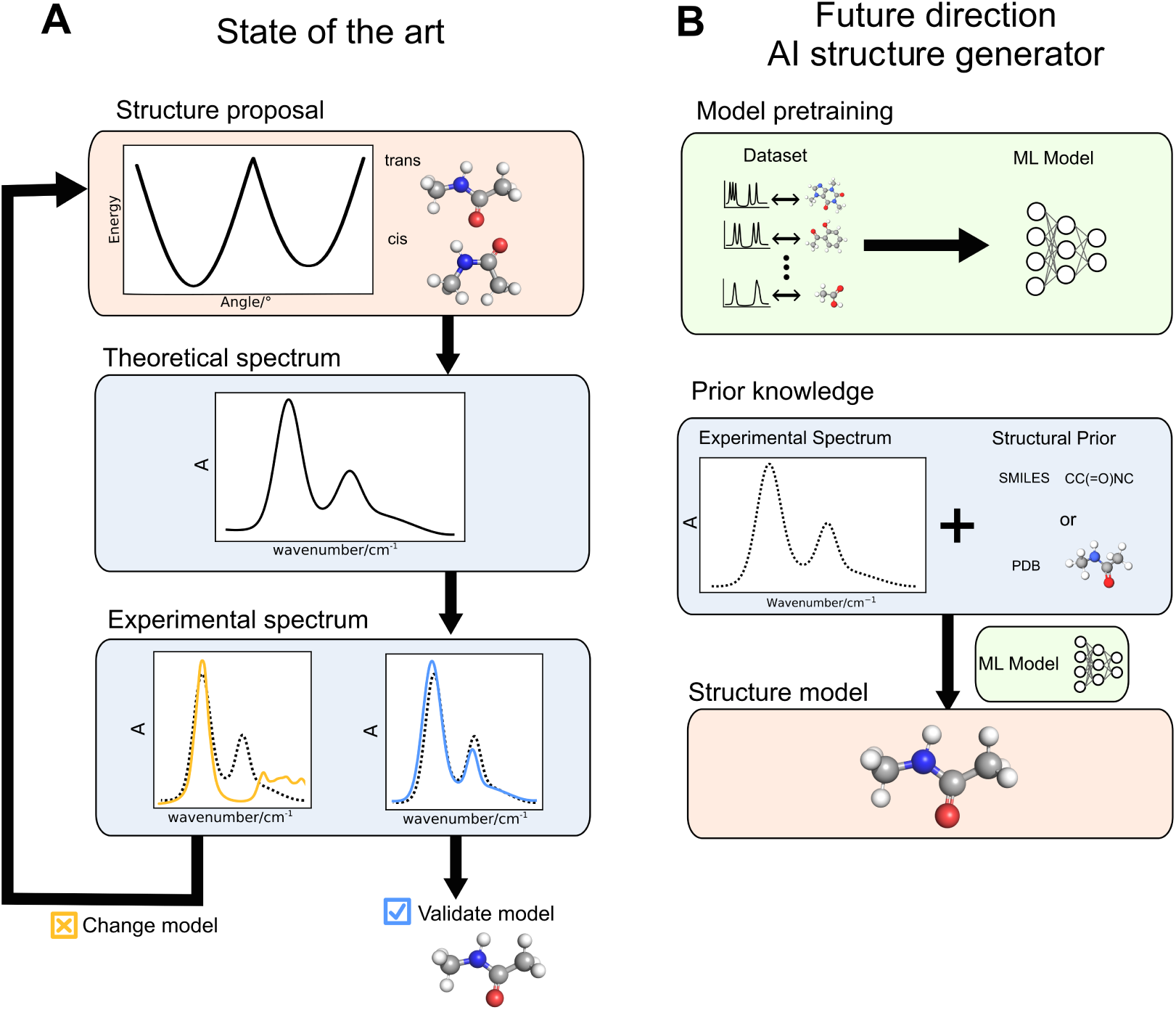
Traditional and ML-enhanced structural spectroscopy strategies. The traditional strategy (A), addressing the forward problem in IR spectroscopy, starts with a structural proposal and a human-in-the-loop iteratively updates the structure by comparing its theoretical spectrum with an experimental reference, until close alignment. The ML-enhanced strategy tackling the inverse problem (B) requires first training a model on a paired spectrum-structure dataset. The experimental spectrum, often conditioned with a structural prior (such as a SMILES string or reference geometry), is then passed as input to the trained model. The model output is a direct prediction of the molecule’s 3D structure based on the experimental spectrum. Other paradigms for ML-enhanced workflows are also possible; see the main text for details.

Increasingly, machine learning (ML) methods are providing new ways of accelerating the forward problem. For instance, ML force fields and dipole models can be trained on density functional theory (DFT) data, enabling MD simulations at a level approaching DFT accuracy but at a fraction of the computational cost, which can then be used with NMA or FT analysis to generate the spectrum.^35^ While accurate, this simulation-based approach is not scalable, as it requires fresh computation for each new molecule. An alternative approach uses supervised ML models to directly map molecular structures to their IR spectra. While early work focused on specific bands like the amide I frequency in proteins,^36^ recent graph neural networks generate full spectra.^37,38^ These models show high correlation (Spearman ∼ 0.9) with ground truth data, but their performance drops significantly for molecules with novel structural features outside of the training data, and there remains substantial room for improvement in predicting absorption intensities and bandwidths.

In addition to accelerating the forward problem, ML techniques are starting to introduce a paradigm shift by attempting to directly solve the *inverse problem*: predicting a molecular structure directly from its spectrum (Figure 1B). This strategy requires training models on large datasets of paired structures and spectra. Although predicting a full 3D structure from an IR spectrum alone is currently an unsolved challenge,^39^ significant progress has been made on simplified tasks like predicting functional groups^40^ or generating molecular graphs.^41^ Accuracy is further improved by incorporating prior knowledge or complementary data, with the integration of NMR spectra proving particularly effective.^42^

Here, we explore how machine learning (ML) can advance IR spectroscopy into a method for analyzing molecular structure and dynamics, comparable to NMR spectroscopy. To assess this potential, we compare six theoretical workflows for generating IR spectra, combining two calculation methods (Normal Mode Analysis and Dipole Autocorrelation) with three MD simulation approaches (Classical MM, QM/MM, and ML-based potentials).

We apply these workflows to N-Methylacetamide, a common benchmark molecule that models a peptide backbone and exists in two distinct conformations, *trans* and *cis*.^43–48^ By validating the calculated spectra against experimental data, we provide a basis for discussing the merits of each simulation and calculation method.

Ultimately, we anticipate that this rigorous integration of experimental and theoretical IR spectroscopy has the potential to become a cornerstone of 4D structural biology, complementing time-resolved X-ray crystallography and cryo-EM. Beyond its broad applicability, this approach holds particular promise for resolving challenging targets such as the heterogeneous aggregates implicated in neurodegenerative diseases such as Alzheimer’s and Parkinson’s. The detailed structural insight gained could lay the foundation for advances in diagnostics and targeted therapies—an achievement that has so far remained out of reach for existing structural biology methods.

## Methods

To account for dynamic ensembles found in IR spectra we make use of biomolecular simulation strategies to generate these ensembles. We compare different simulation approaches using quantum chemical calculations and molecular dynamics (MD) simulations to assess their ability to produce high-quality theoretical IR spectra. The overall workflow is presented in Figure 2 and encompasses the following steps. First, a classical molecular mechanics (MM) simulation was used to equilibrate a solvated starting structure of *trans*- and *cis*-N-Methylacetamide (*cis*-NMA), followed by three different types of biomolecular simulation production runs from which IR spectra are derived. In the following, we present the details of the different methods used to simulate the structural ensemble and to extract the calculated theoretical IR spectra.

**Figure 2:**
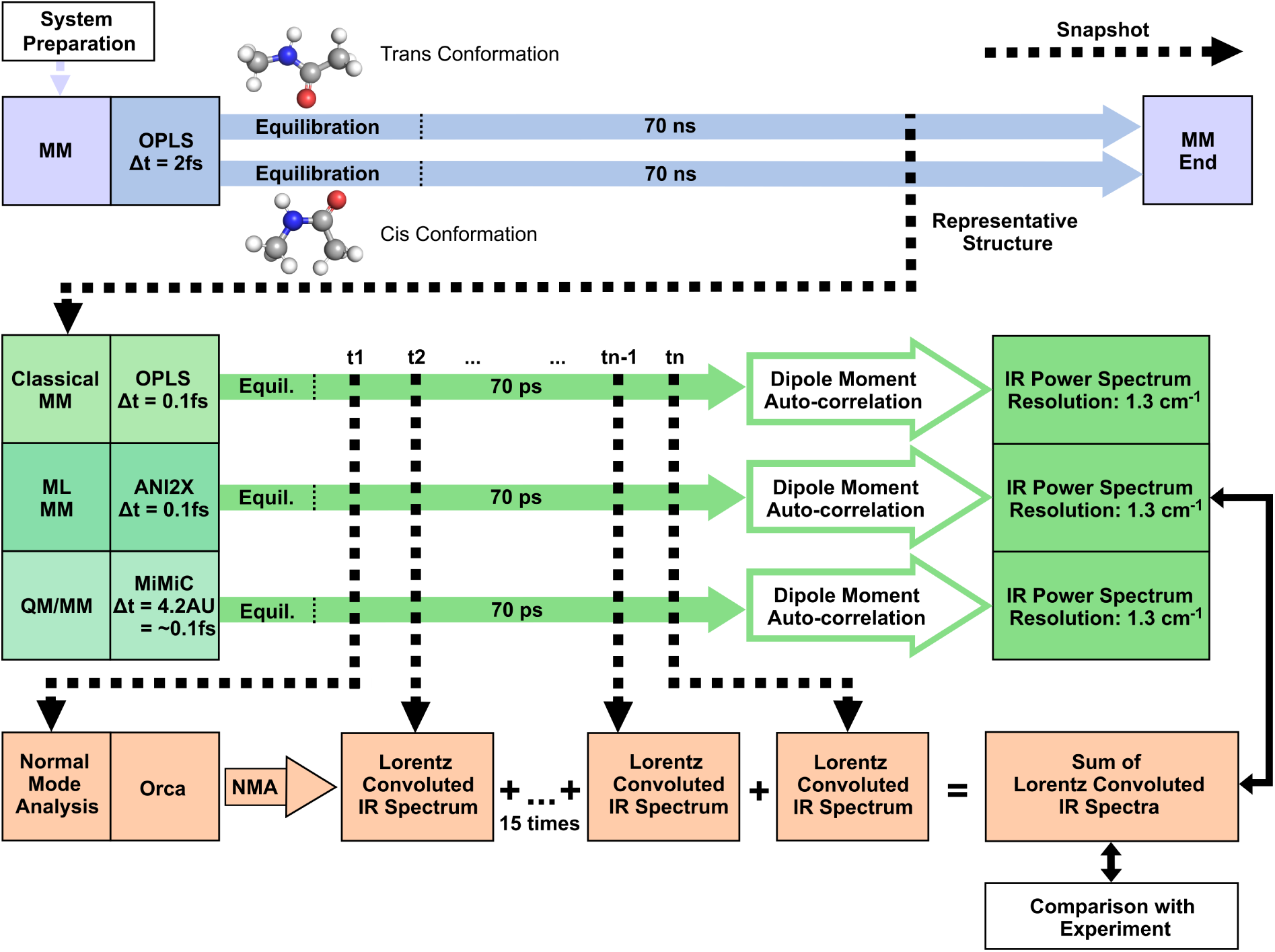
Traditional theoretical IR spectroscopy workflow. First, we performed 70 ns classical molecular mechanics (MM) simulations with a 2 fs step size, to equilibrate the *trans* and *cis* conformations of N-Methylacetamide in solution. We then used the geometries of the representative simulation structure as starting points for the three simulations sampling the atomic motion within one conformation with a step size of 0.1 fs: A classical MM simulation with the OPLS/AA force field,^49^ a machine-learned potential-based MD simulation with ANI-2x^50^ and a QM/MM simulation using MiMiC.^51,52^ From each of these runs, a theoretical IR spectrum was calculated using both NMA and DMA. For validation, these spectra were compared with the experimental IR spectrum. This workflow allows a systematic evaluation of how both the IR calculation method and the underlying simulation technique affect spectral accuracy and conformational differentiation.

### Simulation system preparation

The workflow is initiated by preparing the structures of *trans*-NMA and *cis*-NMA (Figure 2, top left corner) using the software packages MAXIMOBY (v. 2025)^53^ and GROMACS (v. 2024.1).^54,55^ Initially, the first and second solvation sphere were added using the solvation approach implemented in MAXIMOBY that is based on the Vedani algorithm.^56^ To prevent self-interactions due to periodic boundary conditions in the MM simulation, the N-Methylacetamide and its solvation shells were placed in a cubic simulation cell with a minimal distance of 13 Å to the solvens molecules. This resulted in a cubic box with an edge length of 44 Å for *trans*-NMA and 42 Å for *cis*-NMA. The box was then filled with bulk water containing 2754 and 2549 water molecules respectively, using the solvation strategy implemented in GROMACS. To resolve steric clashes in the transition between the second solvation shell and the bulk water, an energy optimization of water hydrogen atoms was performed in MAXIMOBY using the implemented Amber84^57^ united atom force field and the force field corresponding TIP3P water model.^58^ Finally we convert the TIP3P water model to TIP4P.^58^ TIP4P is the corresponding water model for the Optimized Potentials for Liquid Simulations - All Atoms (OPLS/AA) force field^49^ that is used for the subsequent MM simulations. It shows better agreement with experimental measurements for dynamic properties.^58^

### Molecular mechanics (MM) simulations

The prepared structure serves as the starting point for the two initial *trans*-N-Methylacetamide (*trans*-NMA) and *cis*-N-Methylacetamide (*cis*-NMA) MM simulations, illustrated in Figure 2 as two broad, blue arrows. The systems were heated to 293 K during a 1 ns nVT equilibration with a step size of 1 fs. The temperature was controlled using a velocity-rescaling thermostat^59^ with a coupling constant of 0.1 ps. The heating process was carried out in two stages: Initially, the temperature was gradually increased from 0 K to 100 K over the first 100 ps, followed by a further increase to 293 K over the subsequent 900 ps. This temperature was chosen to match the room temperature during the experimental measurements. The system was then equilibrated using an additional 1 ns nVT simulation (step size 1 fs) with a constant temperature of 293 K, followed by a 10 ns npT simulation (step size 1 fs). The temperature coupling is done with the velocity-rescaling thermostat (coupling constant 0.1 ps), and the pressure coupling using a Berendsen barostat^60^ (coupling constant 0.1 ps). Subsequently, a 70 ns npT production run with a step size of 2 fs was performed, using the Nośe-Hoover thermostat^61,62^ (coupling constant 0.5 ps) and the Parrinello-Rahman barostat^63^ (coupling constant 2.5 ps). To allow for a step size of 2 fs, the heavy atom-hydrogen bonds were constrained with the LINCS algorithm.^64^ We extracted a representative structure (see Figure 3) from each of the *trans* and *cis*-NMA production runs to initiate the more detailed MD simulations. The final MM runs (Figure 2, top filled green arrow) were used to extract spectra with NMA and DMA apply the same simulation strategies using a timestep of 0.1 fs and no constraints on any bonds.

**Figure 3:**
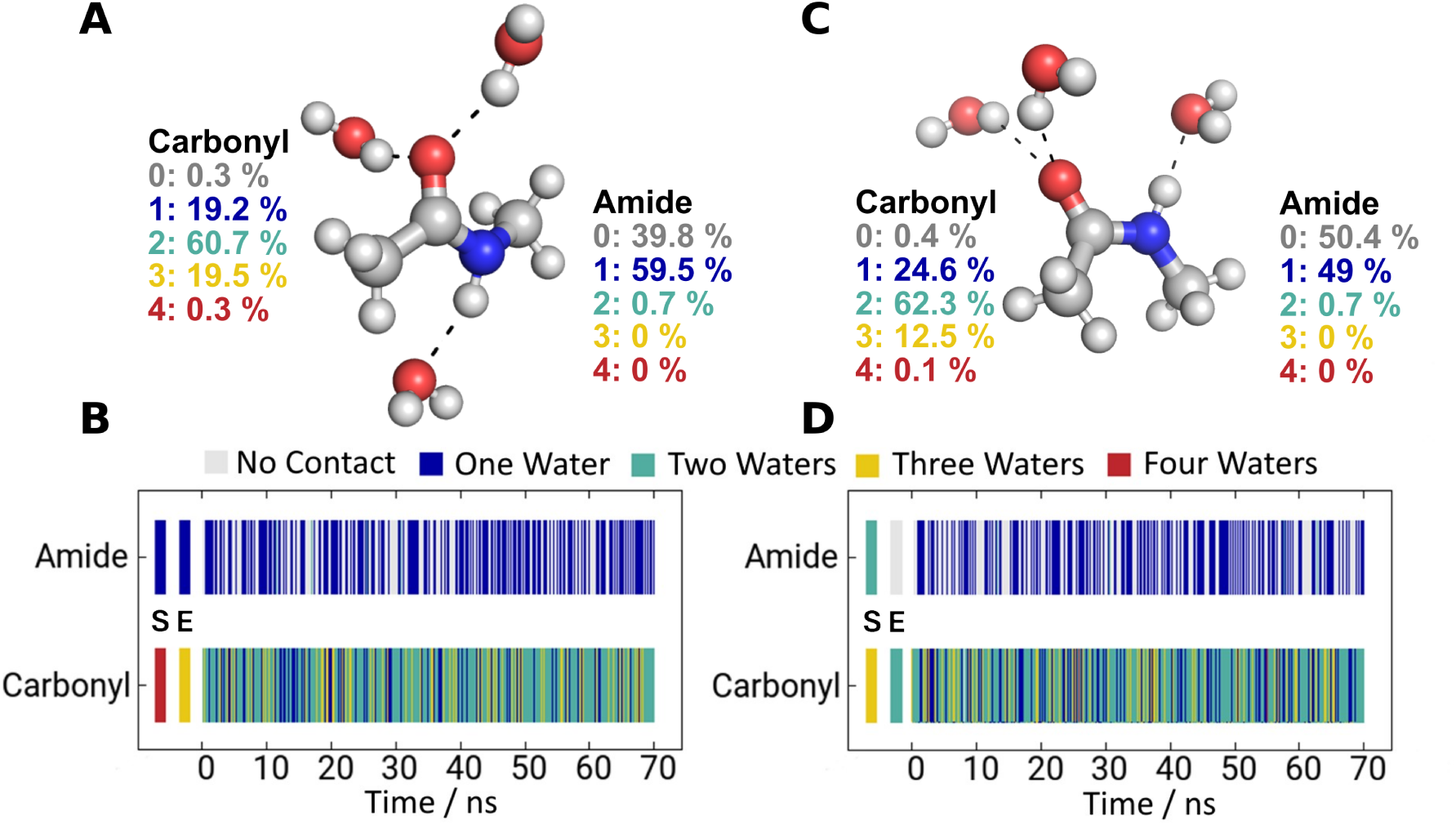
Water interactions with N-Methylacetamide during a 70 ns (2 fs step size) molecular dynamics simulation. Representative structures for the (A) *trans* and (C) *cis*-conformations are shown with direct interacting water molecules; bulk solvent is omitted for clarity. The number of water molecules bound to the carbonyl and amide groups was tracked over the simulation time. On average, two water molecules are bound to the carbonyl and one to the amide for both conformations. The percentage of occupation of different numbers of water molecules over the simulation time is shown next to the structures. Panels (B) and (D) show the contact analysis over time, color-coded for the number of different water molecules, for *trans-* and *cis*-conformations, respectively. The first and second data points, labeled S and E correspond to the starting structure and the structure after heating and equlibration.

### Molecular mechanics (MM) simulation analysis

Inter-molecule interactions between N-Methylacetamide and water molecules were analyzed using PyContact14^65^ and the contact matrix algorithm implemented in MAXIMOBY.^53^ N-Methylacetamide posseses no major internal degrees of freedom apart from the *trans*/*cis* isomerization. The representative structure of the first MM MD simulation was therefore defined by its contacts with the surrounding water molecules. We identified the number of water molecules forming a hydrogen bond to each functionality of N-Methylacetamide every 0.1 ns. On average, about two waters are present at the carbonyl and 1 water at the amide (Figure 3). This was true for both conformations. As there were many structures fulfilling these criteria, we picked the last geometries of the respective simulations doing so and used them as starting structures for the finer MD runs. In Figure 2, these simulation runs are shown as green, broad arrows, originating from green boxes. The three simulation types are listed vertically.

### Molecular dynamics simulations using machine-learned interatomic potentials

Hybrid ML/MD simulations (Figure 2, middle filled green arrow) were initiated by the two representative N-Methylacetamide structures using for each two types of machine-learned interatomic potentials: ANI-2x and MACE-OFF23 (small).^50,66^ ANI-2x, which currently supports C, O, H, N, F, Cl and S atoms, was trained on 8.9 million molecular conformations of drug-like molecules from GDB-11 and ChEMBL.^67,68^ The potential is constructed from atomic environment vectors ^69^ and trained to reproduce molecular energies and forces at the *ω*B97X/6-31G(d) level of DFT theory. MACE-OFF23 supports C, O, H, N, F, Cl, S, P, Br and I atoms and is trained on the SPICE dataset^70^ with an equivariant MACE architecture neural network.^71^ MACE-OFF was trained to reproduce the energies and forces computed at the *ω*B97M-D3(BJ)/def2-TZVPPD level of DFT theory.

In all simulations, the ML potential (either ANI-2x or MACE-OFF23) was used to model forces between atoms of N-Methylacetamide. The molecule was solvated in a water box of classical TIP-4P.^58^ All forces between the water and N-Methylacetamide were modelled with the OPLS/AA force field. ^49^ Simulations were performed in OpenMM 8.2 in the nVT ensemble using the recommended BAOAB-Langevin integrator at 293 K (Langevin thermostat) with a friction coefficient of 1 ps.^72^ The equilibration run was 15 ps with a 1 fs timestep. The production run was 70 ps with a 0.1 fs timestep. Code for reproducing these experiments is available at https://github.com/dominicp6/mlmd.

### Hybrid quantum mechanics/molecular mechanics (QM/MM) simulations

QM/MM calculations (Figure 2, bottom filled green arrow) were performed through the MiMiC interface.^51,52^ MiMiC is a framework to perform multiscale simulations in which loosely coupled external programs describe individual subsystems at different resolutions and levels of theory, particularly suitable for HPC setups.^73^ In this work, MiMiC coupled the DFT-based quantum code CPMD^74^ with the popular classical molecular dynamics code GROMACS.^54,55^ The QM/MM simulations divided the systems into a QM region, which includes the N-Methylacetamide, and an MM part consisting of all water molecules. The QM regions were treated at DFT level of theory with the PBE recipe for the exchange-correlation functional.^75^ The wave function of the QM region was expanded in a plane-wave basis set up to an energy cutoff of 70 Ry. Only valence electrons were explicitly treated, while core electrons were described using norm-conserving pseudopotentials of the Martins-Troullier type.^76^ The MM water molecules were described by the TIP4P model^58^ and the OPLS/AA force field. ^49^

### Dipole moment autocorrelation analysis (DMA)

One method for calculating IR spectra is based on the autocorrelation analysis of the dipole moment. This approach takes advantage of a key principle of infrared spectroscopy: IR light is only absorbed if the molecular vibrations lead to a change in the dipole moment. The dipole moment autocorrelation function captures these fluctuations by monitoring how the dipole moment evolves over time during a molecular dynamics simulation with time steps of 0.1 fs. The DMA is illustrated in Figure 2 following the path of the three MD simulations and resulting in the IR Power Spectrum (green boxes on the right). The dipole moment is calculated by CPMD for every step employing the PBE functional^75^ and a plane wave basis set. To ensure that relevant vibrational modes are sampled adequately, the simulation must run long enough to allow multiple oscillation cycles. The resulting IR power spectrum is obtained by applying a fast Fourier transform (FFT) to the autocorrelation data and subsequently corrected using a quantum correction factor. In molecular dynamics simulations—including QM/MM and CPMD, the motion of atoms is treated classically. As a result, quantum effects in vibrations, especially in fast bond oscillations, are not fully captured. Thus, a quantum correction factor (QCF) is needed to include these effects in the calculated IR spectrum.^77,78^ The here used method provides a spectral resolution of 0.24 cm^-1^, but does not allow a straightforward assignment of individual normal modes.

### Normal Mode Analysis

An alternative method to compute IR spectra is Normal Mode Analysis (NMA). To calculate IR spectra, we performed NMA calculations initiated by structures from each of the three different detailed simulation trajectories (each 70 ps) with 5 ps distance (15 structures). These structures are depicted in Figure 2 as black dotted arrows originating from the three MD simulations. We used the QM/MM embedded NMA frequency calculation method implemented in Orca 5.0.4.^79^ This scheme differentiates between a high layer, consisting of the area that is treated quantum mechanically, and a low layer, consisting of the area that is treated using classical molecular mechanics. The QM part contained the N-Methylacetamide and the closest ten water molecules. We tested varying numbers of QM-treated water molecules. A lower amount yielded a worse experimental agreement, while a higher amount did not improve it and increased computation time. The MM part contained all remaining solvent molecules. The eigenvalues and eigenvectors of the normal vibrations are obtained by diagonalizing the Hessian. One important condition for this calculation is that the structure is in an energetic minimum. To reach a minimum structure we performed an iterative QM/MM and MM optimization in the same way as described by Mann et al.^80^ but replaced the QM software Gaussian by ORCA. First, the partial charges are calculated by ORCA using QM/MM and only the charges are handed over for the MM simulations with MAXIMOBY. Then alternating MM optimizations with MAXIMOBY^53^ and QM/MM optimizations in Orca 5.0.4 were performed three times. The coordinates and charges from each step were passed on to the respective following step. The calculation of the charges in the QM region derived from the electrostatic potential follows the Merz-Singh-Kollman (MK) scheme. The MM optimizations were performed using the Amber5 force field. ^81^ The QM calculations were performed using the functional PBE with the basis set 6-31G*. This turned out to be the most economic functional and basis set combination to calculate IR spectra as shown by an extensive comparison study.^80^ The calculated intensities were fitted with a Lorentz function with a width of 20 wavenumbers and subsequently summed up over the 15 snapshots to generate the final spectrum, illustrated in the orange row in Figure 2.

### Experimental infrared spectroscopy (ATR-FTIR measurements)

ATR-FTIR measurements of N-Methylacetamide in H_2_O were performed using a Vertex-70-spectrometer (Bruker Optik, Ettlingen, Germany) in the double-sided, forward-backward-mode. The spectral resolution was 2 cm*^−^*^1^ and the scanner velocity 16 kHz. The resulting interferograms were processed using the Mertz phase correction, a Blackman-Harris three-term apodization function and a zero filling factor of 4. The ATR-accessory integrated in the spectrometer was the DuraSampl*IR* II (Smiths Detection, London, England) with nine active reflections. For the spectra, an average of 224 scans was used. The background spectra included 112 scans. N-Methylacetamide was obtained from Sigma-Aldrich Chemicals (St.Louis, USA).

## Results and Discussion

Our results and discussion are split into two parts; in the first part, we evaluate different approaches following the traditional strategy of connecting IR spectroscopic data with structural models addressing the forward problem of theoretical IR spectroscopy. In the second part, we discuss how the evaluated approaches pave the way for solving the inverse problem of assigning structures to spectra directly and the potential to advance IR spectroscopy to a direct structure-giving method with sub-Ångström resolution.

### Sampling of the conformational energy landscape results in two stable solvated N-Methylacetamide conformations

N-Methylacetamide is a well-studied^43–48^ benchmark system to model a peptide bond. For IR-spectroscopy to be useful as a structural interrogation method, such benchmark systems are essential to elucidate more complex protein structures in the future. To compare our experimental IR spectrum of N-Methylacetamide to the different theoretical methods, we need a robust understanding of the conformational space of N-Methylacetamide, which we here generated from molecular dynamics simulations. N-Methylacetamide has two dominant conformers *cis*-N-Methylacetamide (*cis*-NMA) and *trans*-N-Methylacetamid (*trans*-NMA), with the *trans* form being 2.5 kcal/mol more stable than the *cis* form.^82–85^ The energy barrier separating these conformers is 18.8 and 21.3 kcal/mol ^86^ respectively, meaning that within even moderate sampling of molecular dynamics trajectory no interconversions between *cis* and *trans* is expected in MD simulations at room temperature in the timescale studied here. As a result, we ran two separate MM simulations of the two conformers to obtain equilibrated, solvated representative structural models of *cis*-NMA and *trans*-NMA (see the blue part of the workflow Figure 2). As expected, no interconversion between *cis* and *trans* isomers is observed in the 70 ns MM trajectories. Going beyond systems with high barriers, different conformers can be identified with sufficiently long MM simulations, and their populations estimated from the simulations. However, in our study, we assigned these two conformations *a priori* and used the initial MM simulation to analyze and identify sub-conformer dynamics and their respective water interactions. The identified representative structures were used to start all production runs for our theoretical IR spectra calculation. The pattern of water molecules forming hydrogen bonds with N-Methylacetamide is the primary criterion for selecting the representative structure of each conformation (Figure 3 A and C).

In the *trans* conformation, two water molecules are hydrogen-bonded to the carbonyl functionality for the majority of the simulation. However, transition states with one or three bound water molecules are frequently observed. The amide has the tendency to form only one hydrogen bond. One water molecule is present in ∼60 % of the simulation time, while the remaining ∼40 % corresponds to transition states without a clearly defined hydrogen bond. The *cis* conformation shows the same general trend, albeit with a slight shift to a lower average number of hydrogen bonds, while the median remains unchanged. The representative structures used as starting points for the following evaluation of theoretical IR spectroscopy approaches were chosen to reflect these hydrogen bonding patterns and are shown in Figure 3 A for *trans* and C for *cis*, respectively, while the frame-wise contact analysis is shown in Figure 3B and D.

### Efficient workflows for evaluating traditional theoretical IR spectroscopy enable the connection between structural and spectral data

We calculated and measured IR spectra of N-Methylacetamide to evaluate the different strategies that connect IR spectroscopic data with structural models addressing the forward problem (Figure 1 A). The experimentally measured spectrum serves as a gold standard to evaluate the calculation strategies. An overview of the IR spectra calculation workflow is illustrated in Figure 2. Calculation parameters and experimental conditions are detailed in the Methods section. The initially determined representative structures of the conformational energy landscape minima are used to investigate the detailed atomic motion within a given N-Methylacetamide conformation. We performed MD simulations using three different levels of theory: classical molecular mechanics (MM) forcefield-based, machine-learned potential-based molecular dynamics (ML), and hybrid quantum mechanics/molecular mechanics (QM/MM) simulations. To calculate the IR spectra based on the structural data, we used the two state-of-the-art approaches: normal mode analysis (NMA) and dipole moment autocorrelation (DMA).

The resulting spectral comparison is presented in Figure 4. For spectral analysis, two key spectral regions were considered: (i) the amide I and II bands (1500-1700 cm*^−^*^1^), reflecting the C-O stretching and a combination of C-N stretching and N-H bending vibrations respectively, and (ii) the fingerprint region (1250-1450 cm*^−^*^1^) which includes methyl-bending and amide III modes. The peak positions in these regions are listed in Table 1, while Table 2 summarizes the deviations between calculated and experimental peak positions. Since assigning atom groups to each peak using DMA is highly challenging and controversial, we focus instead on the peak pattern defined by the spectral band positions and intensities.

**Figure 4:**
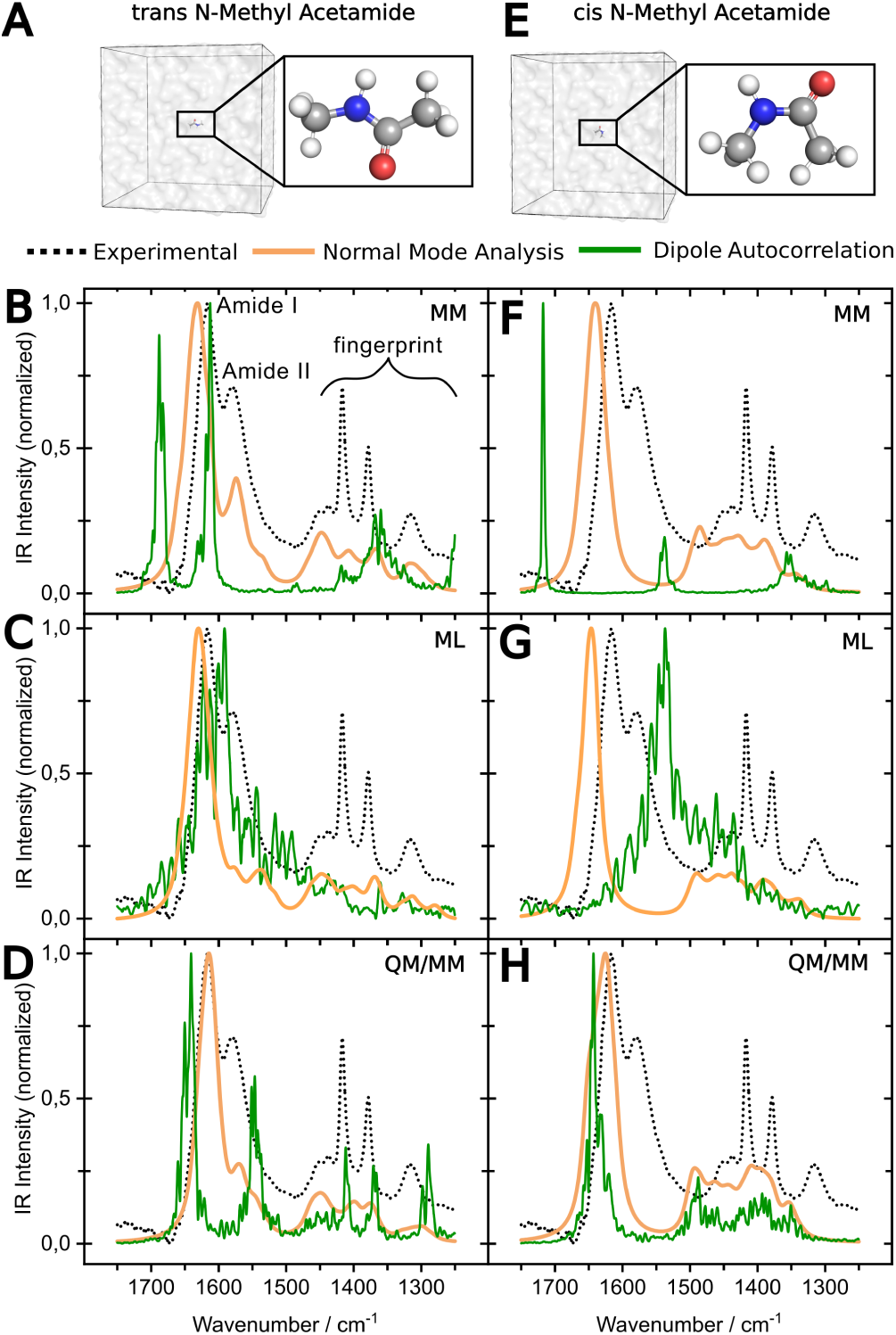
Comparison of theoretical and experimental IR spectra. **A** shows the simulation system for solvated *trans*-NMA and **E** the one for *cis*-NMA. The left column shows the theoretical IR spectra calculated based on NMA (orange) and dipole moment auto-correlation (green) for *trans*-NMA (**B-D**) and the right one for *cis*-NMA (**F-H**) compared to the experimental spectrum (black dashed line). Compared are three different methods to obtain the input geometries for the spectra calculation, namely MM (**B,F**), machine learned MD (ANI-2x results shown) (**C,G**), and QM/MM using MiMiC (**D,H**).

**Table 1:**
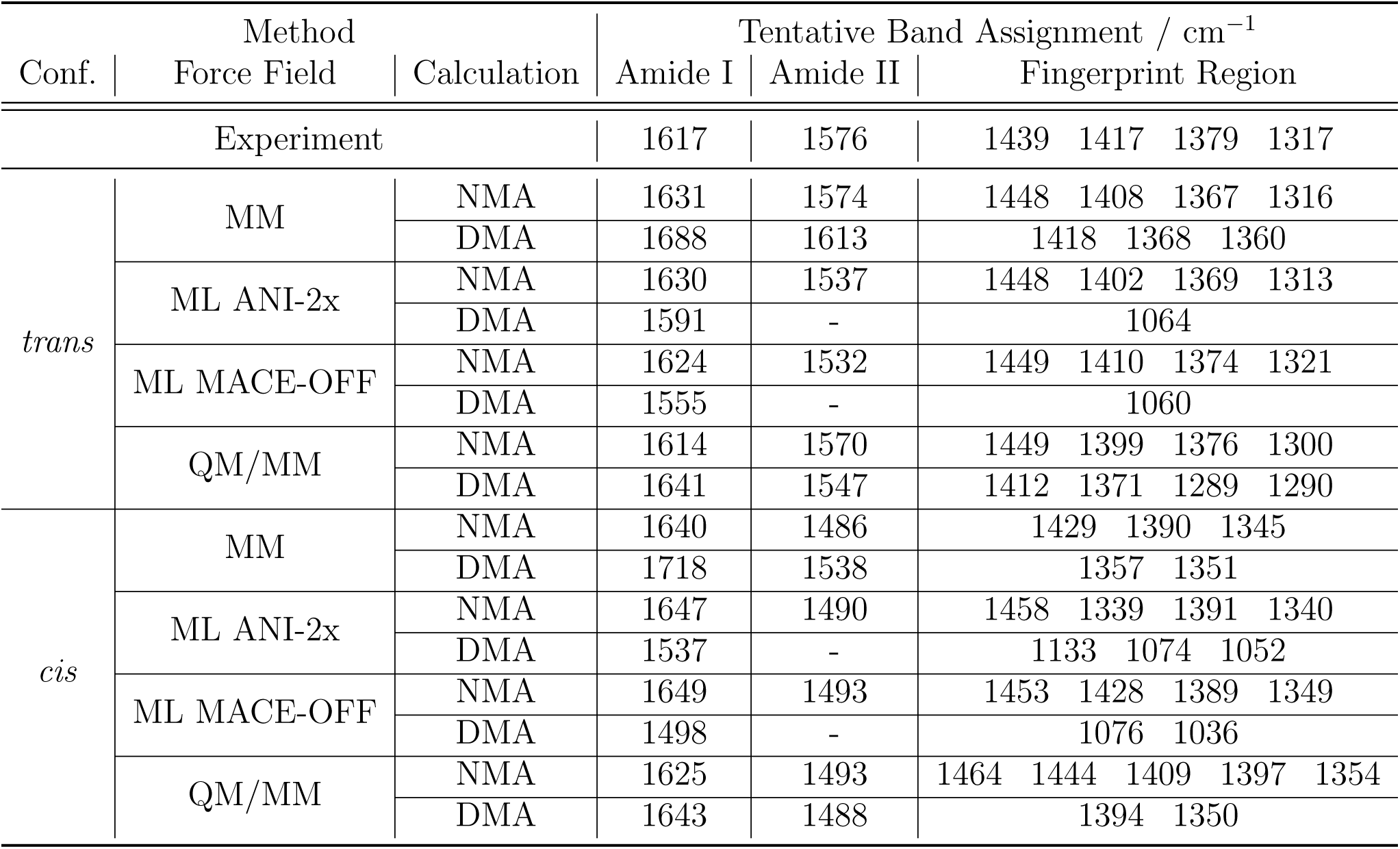
Comparison of the calculated and experimental peak positions.

**Table 2:** Deviation of the assigned calculated peak positions to the experimental ones.

| Conf. | Method | | Deviation from Experiment / $cm^{-1}$ | | Intensity Ratio |
| --- | --- | --- | --- | --- | --- |
|  | Force Field | Calculation | Amide I | Amide II | Amide I / Amide II |
| Experiment |  |  | 0 | 0 | 1.40 |
| *trans* | MM | NMA | +14 | -2 | 2.52 |
|  |  | DMA | +71 | +37 | 0.89 |
|  | ML ANI-2x | NMA | +13 | -39 | 5.92 |
|  |  | DMA | -26 | - | - |
|  | ML MACE-OFF | NMA | +7 | -44 | 7.14 |
|  |  | DMA | -62 | - | - |
|  | QM/MM | NMA | -3 | -6 | 3.63 |
|  |  | DMA | +24 | -29 | 1.79 |
| *cis* | MM | NMA | +23 | -90 | 4.36 |
|  |  | DMA | +101 | -38 | 5.16 |
|  | ML ANI-2x | NMA | +30 | -86 | 6.40 |
|  |  | DMA | -80 | - | - |
|  | ML MACE-OFF | NMA | +32 | -83 | 6.05 |
|  |  | DMA | -119 | - | - |
|  | QM/MM | NMA | +8 | -83 | 3.83 |
|  |  | DMA | +26 | -88 | 4.36 |

### All combinations of theoretical methods clearly distinguish between the *trans* and *cis* conformations, with all *trans*-NMA spectra showing markedly better agreement with the experimental data than their *cis* counterparts

All *cis*-NMA spectra calculated by NMA consistently exhibit two bands in the amide region that are separated from each other. While the dominant calculated peak is in the region of the amide I, the second peak is red-shifted to the region around 1485 cm*^−^*^1^. This strongly red-shifted second peak is not present in the experiment and represents the key marker band to distinguish between *trans*- and *cis*-NMA. This clear peak separation is also observed in the calculated spectra from DMA based on the MM and QM/MM data. In contrast, the ML-based DMA spectra for both *cis*- and *trans*-NMA show only one dominant peak with a broad shoulder. However, the wavenumber of this maximum peak strongly differs between the conformations allowing us to distinguish them. The *cis*-NMA peak is red-shifted by ∼54 cm*^−^*^1^ compared to the one of *trans*-NMA using ANI-2x and ∼57 cm*^−^*^1^ using MACE-OFF (Table 1).

By thermodynamical preference, N-Methylacetamide exists predominantly in the *trans* conformation (98.5 %), as *trans*-NMA is 2.5 kcal/mol more stable than *cis*-NMA.^87,88^ Under these thermodynamic considerations, we find that all the calculated spectra for the *trans* conformation agree better with the experiment than the calculated spectra for the *cis* conformation. In systems with more equally distributed conformations under experimental conditions, population-weighted IR spectra need to be calculated using MM simulations or other methods to sample the conformational space. Such weighting has been successfully applied to flexible molecules in solution, demonstrating the importance of thermodynamic sampling in spectral predictions.^89^ In the case of N-Methylacetamide, however, the overriding dominance of *trans*-NMA (98.5%) makes the need for population-weighting negligible. We consider only the results for *trans*-NMA for our evaluation in the following.

#### All *trans*-NMA spectra calculated by Normal Mode Analysis represent the experimental features well, with the best agreement for the QM/MM-based method

All simulation approaches using NMA as the subsequent spectra calculation method successfully reproduced the main spectral characteristics of *trans*-NMA observed experimentally. These include a prominent amide I band, a detectable amide II band, and characteristic bands in the fingerprint region. However, there are some detailed deviations from the experimental spectrum:

Firstly, for all methods, deviations are observed in the relative band intensities. Specifically, the calculated amide I/II ratio is not in agreement with the experimental data, due to an underestimation of the amide II intensity. Moreover, the calculated peak around 1450 cm*^−^*^1^ is always the most prominent one in the fingerprint region, while it is the weakest in the experimental data. The remaining fingerprint bands are broadened and less intense than experimentally observed.

Secondly, a detailed analysis of the amide region reveals that for both the MM and the QM/MM-based spectra, the amide I and II peaks are in very good agreement with the experiment. In contrast, the ML-based spectrum does not reflect the experimental pattern as the amide II peak is red-shifted with low intensity. The spectra of the two ML interatomic potentials ANI-2x and MACE-OFF display an identical general shape and share all major spectral characteristics, with only slight deviations regarding the peak positions. Figure 4 displays the results using ANI-2x. A comparison between the two ML-based potentials is found in Supporting Figure S1 and S2.

Thirdly, the number of calculated peaks and their positions within the fingerprint region agree with the experiment for all methods. However, the overall pattern is not reproduced in detail by any of the methods, as the intensity and band broadening of each of the four peaks deviates from the experiment.

In summary, the MM and QM/MM-based NMA calculated peak positions agree with the experimental data, but further refinement of the method is needed to improve the amide I and amide II intensity ratios and the shape of the fingerprint region. The QM/MM-based result is slightly better than the MM-based one, which in turn is in better agrement than the ML-based result. The very similar results of MM and QM/MM-based spectra indicate that the OPLS/AA MM force field accuracy is close to the QM accuracy, and QM optimization of snapshots is already sufficient to capture the key spectral features. This holds true for at least simple structures with limited dynamics within a conformation.

### Examining the Dipole Moment Autocorrelation-based calculations, the QM/MM-derived spectrum matches the *trans*-NMA experimental features best

The DMA approach depends even more strongly on the accuracy of the underlying MD simulation approach than the NMA. For DMA, every written structure (all 700,000 frames at 0.1 fs) of the simulation accounts for the calculation, while for NMA, only 15 QM optimized structures are used. Therefore, the deviation between the three MD methods is stronger than for the NMA calculated spectra.

Among all DMA calculated spectra, the experimental overall pattern in the amide region is best described by the ML-based ANI-2x potential spectrum. The high intensity peak is red-shifted by 26cm*^−^*^1^ compared to the experiment. However, for the ML-based MACE-OFF potential (Supporting Figure S1) the high-intensity peak is red-shifted by 62cm*^−^*^1^ (Table 2) and broadened relative to the experiment. In contrast, the MM and QM/MM-based spectra show two distinct, well-separated peaks, not in accordance with the experimental shape. The position of the tentativ amide I peak deviates with 26cm*^−^*^1^ in a comparable magnitude to the ANI-2x spectrum, but is blue-shifted in contrast to it. The MM Peak deviates most clearly from the experiment with a blue-shift of 71cm*^−^*^1^. Interestingly, the MM and QM/MM-based spectra both almost exactly reproduces the amide I and amide II intensity ratios (Table 2). The fingerprint region reveals the most pronounced differences between the different approaches. In both ML-based spectra, this region is red-shifted by approximately 300 cm*^−^*^1^ and lacks a well-defined peak pattern (Supporting Figure S2). The MM-based spectrum reproduces three of the four experimental peaks (Table 1). The QM/MM-based spectrum shows four peaks in close proximity to the four experimental ones and overall exhibits the best agreement with the experiment, although deviations remain.

### Among all methods, the QM/MM-based NMA spectrum shows the best agreement with the experiment, although further refinement should be considered

The results demonstrate that the choice of simulation method and IR calculation technique strongly influences the agreement with experimental IR spectra and even the ability to distinguish conformers. QM accuracy is needed to reproduce experimental data, as indicated by the best experimental fit of the QM/MM-based spectra among all DMA calculations. However, MM simulations with subsequent QM/MM optimization provides the most economic way to obtain a solid agreement with the experiment compared to the needed computational power. The shifted fingerprint region for the ANI-2x and MACE ML interatomic potential indicates a clear issue with the way the methyl parameters are learned, suggesting that improvements are needed. Nevertheless, we anticipate that ML-based force fields are the future and will advance theoretical IR spectroscopy methods. However, to date, there is still a clear need to improve such force fields to reach QM accuracy with less computational costs.

Among all methods, the best agreement between calculated and experimental spectra is observed for QM/MM-based NMA calculations. Nevertheless, further refinement remains possible, especially for the peak pattern of the fingerprint region. This region is, in fact, best reproduced by the QM/MM-based DMA calculations. All in all, the differences in peak positions and shapes across methods reflect the sensitivity of the calculated spectra to the quality of detailed atomic changes. This sensitivity underlines the high potential of IR spectroscopy to advance as a crucial structure-giving method by combining theoretical and experimental IR spectroscopy.

### Reliable methods for computational predicting IR spectra pave the way to solve the inverse problem of assigning structures to spectra directly

The power of traditional theoretical IR spectroscopy approaches is evident, nevertheless, the applicability of these traditional approaches becomes increasingly constrained as molecular systems grow in size and complexity. In particular, systems involving the explicit consideration of surroundings, such as proteins or condensed-phase environments, pose significant challenges. The computational expense of quantum mechanical calculations scales poorly with system size, and the harmonic approximation inherent in NMA often fails to capture anharmonic effects and environmental influences that are critical in such contexts. Although MD-based approaches incorporate some of these effects, they remain time-consuming, computationally intensive, and require extensive sampling, particularly for large or flexible systems, where achieving sufficient sampling to capture all relevant conformations becomes increasingly challenging. Moreover, the outcomes of MD simulations can strongly depend on the chosen starting structure or the employed parameters, which may limit the exploration of the conformational space.

The ideal instead is, from just an experimental spectrum as input, to determine conformers and vibrational modes that give rise to different parts of the spectrum without biomolecular simulations or other computationally intensive modeling approaches. This is also known as the inverse problem.^39^

While, to date, we cannot reliably solve for this inverse problem, recent advances in machine learning (ML), particularly in deep learning, offer opportunities to streamline the workflow and possibly increase the accuracy of the mapping between conformers and their vibrations, capturing a specific spectrum of interest. For example, ML-derived force fields accelerate dynamical modeling, substituting the role of QM/MM or DFT approaches ^90^ are a promising alternative, but still leave room for improvement, as demonstrated by the poor capturing of the methyl vibrations in the fingerprint region of *trans*-NMA. Additionally, ML algorithms automate spectra pre-processing tasks such as denoising, spike removal, baseline correction, and feature extraction.^91–93^ ML also aids in hybrid workflows by assisting in ranking candidate structures prior to further refinement.^94,95^

Progress in the inverse problem itself is ongoing.^39^ Current approaches simplify the problem by predicting molecular graphs or SMILES sequences instead of full 3D structures or focusing on easier tasks such as classifying functional groups. Convolutional neural networks have been employed for functional group prediction using FTIR data,^40^ and models combining IR data with molecular formulae have been used to generate SMILES strings.^41^ Integrating diverse spectral data, particularly NMR, generally outperforms methods relying solely on IR for structure elucidation, although 3D molecular structure prediction still remains a challenge.^42,96–98^ Emerging deep generative models, such as diffusion and flow-matching techniques, show promise in tackling these complex, 3D structural inverse problems.^99^ Flow matching has already been applied to 3D structure elucidation from Raman rotational spectra.^100^

A major challenge is in obtaining suitable datasets for training the models. Common benchmarks such as QM9, which include small molecules up to nine heavy atoms, are often supplemented with simulated spectra which fail to capture the full complexity of real data.^39,93,101^ Another challenge is how to incorporate implicit chemical principles - such as valency rules, ring strain or stereochemical preferences - directly into the learning process. Without these, models can struggle to generalize and scale to larger molecules.^39^ Moreover, while many methods achieve promising top-*k* accuracy (how often the correct structure is found in the *k* most probable predictions), achieving high top-1 accuracy is still difficult.^42^ Although recent literature has explored providing confidence estimates to predictions, ^42^ explaining *why* a prediction was made remains an open challenge.

Looking ahead, machine learning (ML) is poised to transform the theoretical analysis of infrared spectra by enabling capabilities that extend beyond the traditional computational methods. As larger and more diverse datasets of experimental and computed spectra become available, ML models will increasingly be able to capture the intricate relationships between molecular structure and vibrational features.

Such capabilities would significantly enhance our ability to interpret spectral data, particularly in cases where traditional approaches struggle, such as in complex mixtures, flexible biomolecules, or materials under dynamic conditions. Ultimately, this may enable automated structure elucidation and spectral interpretation, with potential applications ranging from high-throughput screening to in situ monitoring of chemical processes.

While quantum mechanical methods will continue to provide the theoretical foundation and interpretative depth essential to vibrational spectroscopy, ML-based techniques are expected to play an increasingly central role. With ongoing advances in model architectures, training strategies, and integration with experimental workflows, the coming years are likely to witness a profound shift toward hybrid approaches that combine the strengths of physics-based and data-driven methods for theoretical IR spectroscopy.

## Conclusion

In summary, we demonstrate that current theoretical computational biophysics approaches, particularly MD simulations in combination with QM/MM optimization and normal mode analysis, accurately reproduce key features of experimental IR spectra and distinguish molecular conformations such as *cis*- and *trans*-NMA. This proves that the combination of experimental and computational IR spectroscopy is capable of mining the structural information encoded in the measured data. The sensitivity of vibrational spectroscopy allows for obtaining structural models with sub-Ångstöm resolution, limited by the accuracy of the calculation method. However, the required iterative feedback loop between computational structure prediction and experimental validation is time-consuming and computationally costly. It becomes particularly challenging when the method is applied to molecules with multiple conformations or larger proteins. However solving this challenge is desirable as it will be essential for, e.g., resolving heterogenous aggregates, such as those implicated in neurodegenerative diseases, which are currently unresolvable by existing structural biology methods. Identifying the structure of such drug targets will assist to improve target therapy and diagnostics in future.

Building on recent advances in artificial intelligence tools, the logical next step is to move beyond these forward models and develop methods that directly infer structural information from experimental IR spectra, the so-called inverse problem. We anticipate that machine learning, which has already proven useful in many related fields, will play a crucial role in this endeavor. Its application to this specific problem is still in its infancy with foundational work needed to establish reliable models, generate diverse training data, and incorporate chemical knowledge. Solving the inverse problem will pave the way for advancing IR spectroscopy as a structure-giving method, providing structural and dynamic models with sub-*Å* resolution even of heterogenous structures.

## Acknowledgement

The authors thank Udo Höweler for code development support and fruitful discussions on the different theoretical IR spectroscopy methods. The presented research was funded by the Center for Protein Diagnostics (PRODI), Ministry of Culture and Science of North-Rhine Westphalia, Germany; the United Kingdom Research and Innovation (grant EP/S02431X/1), UKRI Centre for Doctoral Training in Biomedical AI at the University of Edinburgh, School of Informatics. The authors gratefully acknowledge the Gauss Centre for Supercomputing e.V. (www.gauss-centre.eu) for funding this project by providing computing time through the John von Neumann Institute for Computing (NIC) on the GCS Supercomputer JUPITER — JUWELS^102^ at Jülich Supercomputing Centre (JSC). For the purpose of open access, the author has applied a Creative Commons attribution (CC BY) licence to any author-accepted manuscript version arising.

## Notes

### Competing Interest Statement

The authors have declared no competing interest.

