## Supplemental Figures for "IR spectroscopy: from experimental spectra to high-resolution structural analysis by integrating simulations and machine learning"

### Supporting Information

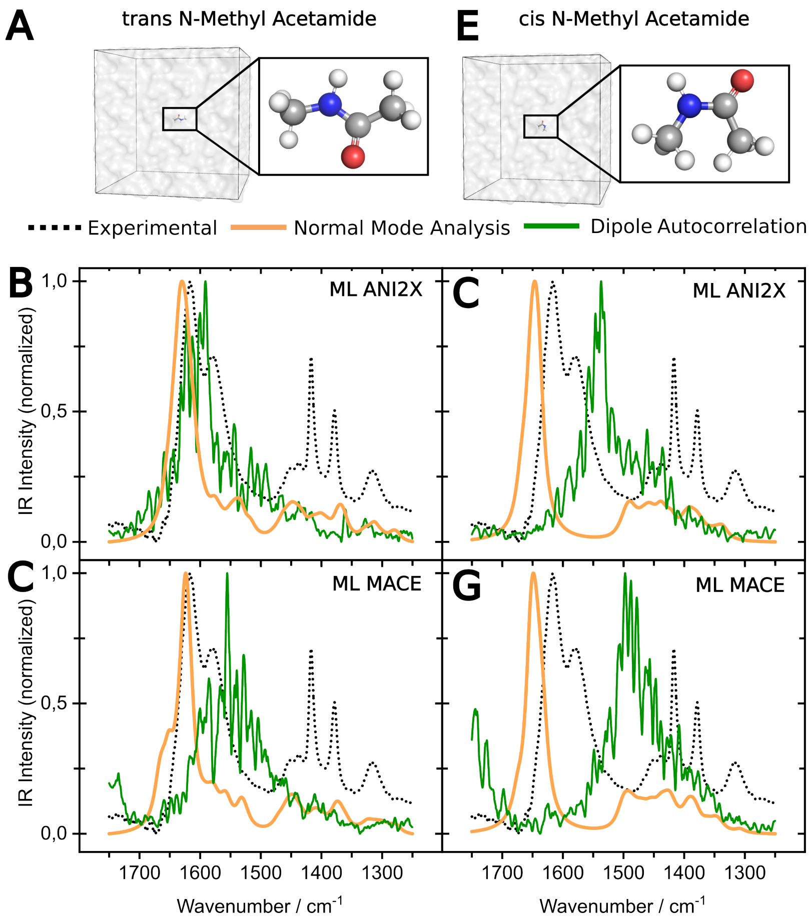


**Figure S1: Comparison of theoretical and experimental IR spectra of N-Methylacetamide.** **A** shows the simulation system for solvated *trans*-NMA and **E** the one for *cis*-NMA. The left column shows the theoretical IR spectra calculated based on NMA (orange) and dipole moment auto-correlation (green) for *trans*-NMA (**B,C**) and the right one for *cis*-NMA (**E,F**) compared to the experimental spectrum (black dashed line). Compared are two different machine-learned force fields used in a simulation to obtain the input geometries for the spectra calculation, namely ANI2X (**B,E**) and MACE (**C,F**).

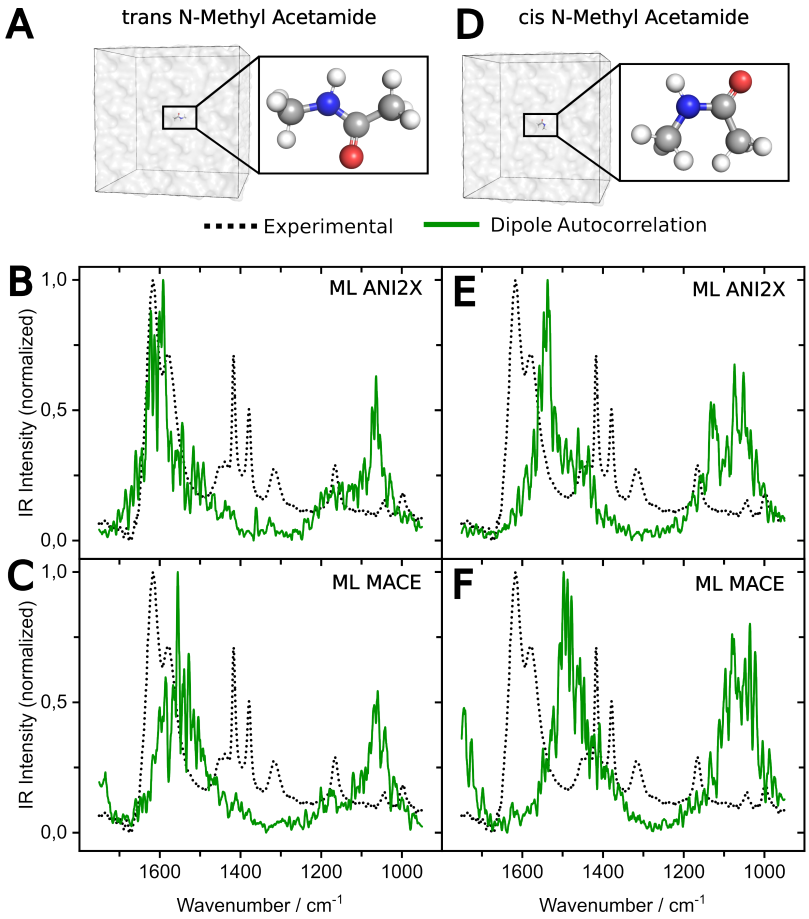


**Figure S2: Comparison of theoretical and experimental IR spectra. A** shows the simulation system for solvated *trans*-NMA and **E** the one for *cis*-NMA. The left column shows the theoretical IR spectra from 950-1450 $cm^{-1}$ calculated based on Dipole Moment Autocorrelation (DMA) for *trans*-NMA (**B**,**C**) and the right one for *cis*-NMA (**E**,**F**) compared to the experimental spectrum (black dashed line). Compared are two different machine-learned force fields used in a simulation to obtain the input geometries for the spectra calculation, namely ANI2X (**B**,**E**) and MACE (**C**,**F**).
